# Pre-trained Maldi Transformers improve MALDI-TOF MS-based prediction

**DOI:** 10.1101/2024.01.18.576189

**Authors:** Gaetan De Waele, Gerben Menschaert, Peter Vandamme, Willem Waegeman

**Affiliations:** Department of Data Analysis and Mathematical Modelling, Ghent University; Laboratory of Microbiology, Ghent University

**Keywords:** MALDI-TOF MS, Neural networks, Transformers

## Abstract

For the last decade, matrix-assisted laser desportion/ionization time-of-flight mass spectrometry (MALDI-TOF MS) has been the reference method for species identification in clinical microbiology. Hampered by a historical lack of open data, machine learning research towards models specifically adapted to MALDI-TOF MS remains in its infancy. Given the growing complexity of available datasets (such as large-scale antimicrobial resistance prediction), a need for models that (1) are specifically designed for MALDI-TOF MS data, and (2) have high representational capacity, presents itself.

Here, we introduce Maldi Transformer, an adaptation of the state-of-the-art transformer architecture to the MALDI-TOF mass spectral domain. We propose the first self-supervised pre-training technique specifically designed for mass spectra. The technique is based on shuffling peaks across spectra, and pre-training the transformer as a peak discriminator. Extensive benchmarks confirm the efficacy of this novel design. The final result is a model exhibiting state-of-the-art (or competitive) performance on downstream prediction tasks. In addition, we show that Maldi Transformer’s identification of noisy spectra may be leveraged towards higher predictive performance.

All code supporting this study is distributed on PyPI and is packaged under: https://github.com/gdewael/maldi-nn

## 1. Introduction

Matrix-assisted laser desportion/ionization time-of-flight mass spectrometry (MALDI-TOF MS) is a proteomic technique commonly used to identify microbial species (Dauwalder et al., 2023). First introduced to clinical microbiology at the end of 2010, its routine use is characterized by low-cost, speed, and reliability (Weis et al., 2020a). MALDI-TOF MS generates spectra containing peaks signifying mostly ribosomal proteins (Seng et al., 2009). As such, the spectra can serve as fingerprints indicative of species identity (Bizzini et al., 2011).

For bacterial species identification, clinical diagnostic labs will typically use the solutions provided by MALDI-TOF MS manufacturers. These solutions are built on large, proprietary, in-house databases (Van Belkum et al., 2012). The models used in such solutions presumably rely on querying certain marker peaks to a large database (Florio et al., 2018). This strategy works reasonably well for identification of most species, but some strains remain problematic to identify this way (Cao et al., 2018; Vrioni et al., 2018). In addition, the peak-matching approach does not suit more-difficult prediction tasks such as strain typing (Hettick et al., 2006), antimicrobial resistance prediction (Weis et al., 2022), and virulence factor detection (Gagnaire et al., 2012). In these cases, researchers turn to machine learning in order to possibly mine more-intricate patterns from the spectra.

Historically, machine learning for MALDI-TOF MS has been hampered by a lack of large open data. Because of this, the nascent field has not often progressed beyond off-the-shelf learning techniques (Weis et al., 2020a). Only a handful of examples exist of more advanced machine learning methods specifically adapted to a MALDI-TOF-based task (Vervier et al., 2015; Weis et al., 2020b; Mortier et al., 2021; De Waele et al., 2023). As such, the design of machine learning algorithms specifically for MALDI-TOF mass spectra has not been studied yet in sufficient detail.

Typically, MALDI-TOF MS-based machine learning methods have either discretized the (*m*/*z*) -axis, or have pre-selected a number of peak locations as features based on training set characteristics. Discretizing the (*m*/*z*) -axis results in fixed-length spectral representations, where every feature consists of a bin. If bins are chosen too large, resolution is lost and multiple peaks may be lumped together. On the other hand, small bins result in a higher-dimensional representation where most bins contain no peaks, unnecessarily adding model complexity (Weis et al., 2020b; Mortier et al., 2021). Other works proposed to select a fixed set of (*m*/*z*) locations based on training set characteristics (Tran et al., 2021). Features for every spectrum are then derived by encoding the peak height (or binary peak presence) for every selected (*m*/*z*) location. The disadvantage here is that spectra may contain important peaks not included in the fixed set of selected (*m*/*z*) locations. Hence, the model is unable to cope with patterns differing too much from general training set characteristics (e.g. rare peaks).

The information in MALDI-TOF mass spectra is defined by their peaks (many associated with ribosomal subunits) (Ryzhov and Fenselau, 2001). As such, we argue that, ideally, a machine learning model for MALDI-TOF mass spectra operates directly on sets of peaks as inputs. This, however, constitutes a non-trivial machine learning setup. A kernel-based technique that processes spectra in this way was previously proposed by Weis et al. (2020b). However, their method is difficult to scale to larger sample sizes. Consequently, in this work, we have developed a deep learning-based solution operating on sets of peaks: a transformer for MALDI-TOF MS data.

Transformers are an ideal neural network candidate for learning on mass spectra for several reasons. Unlike MLPs and CNNs, they do not require a fixed-length feature vector, and hence do not require binning the (*m*/*z*) -axis. Further, unlike RNNs, they do not assume a sequentiality to the signal. Rather, transformers use a permutation-equivariant self-attention mechanism to perform message passing on a complete directed graph (i.e. peaks are nodes, and all nodes are connected to each other) (Vaswani et al., 2017). As such, transformers can be viewed as operating on input sets, rather than sequences.

Because transformers can be viewed as operating on a complete digraph, they place no inductive bias on the learned patterns between input tokens. This property lends transformers supreme representational capabilities, but is also the reason why they are typically described as data-hungry. Consequently, transformers are usually mentioned in the same breath as self-supervised learning (SSL) (Liu et al., 2021). SSL is the paradigm within deep learning wherein a supervised learning task is designed for data that has not explicitly been labeled (Balestriero et al., 2023). This task is (usually) not a useful prediction problem in itself, but rather serves to pre-train a large model. In doing so, greater downstream performance on tasks of interest can be obtained. SSL is, therefore, most useful in scenarios where labeled data is limited, such as the MALDI-TOF MS domain (Weis et al., 2020a).

As related works, following their rise to dominance in natural language processing, transformers are increasingly adopted towards biological data modalities (Clauwaert and Waegeman, 2020; Jumper et al., 2021; Avsec et al., 2021; Elnaggar et al., 2021). Recently, they have also been applied to the mass spectral data domain for (1) *de novo* peptide sequencing (Yilmaz et al., 2022; Yang et al., 2024), and (2) annotating metabolite mass spectra (Goldman et al., 2023; Butler et al., 2023). In the latter data domain, researchers have succesfully used the masked language modeling paradigm (Devlin et al., 2018) to pre-train transformers towards better performance (Bushuiev et al., 2024).

In the present study, we have developed Maldi Transformer, a deep learning architecture for processing MALDI-TOF mass spectra. The inputs to Maldi Transformer consist of spectra as sets of peaks. To fully take advantage of the transformer architecture’s representational capacity, we have proposed a novel self-supervised pre-training strategy. Our strategy relies on discriminating real peaks from noisy ones introduced to the spectrum via shuffling peaks across samples. We pre-trained Maldi Transformer on the large open DRIAMS database (Weis et al., 2020a). Maldi Transformer obtains strong performance on downstream benchmarks, demonstrating the power of the approach.

## 2. Methods

To train and test models, we use three large (> 10 000 spectra) MALDI-TOF datasets, two of which are available to the public domain (§2.1). We define a transformer for MALDI-TOF MS data, and design a novel pre-training task (§2.2). We experimentally validate the final architecture on a number of downstream tasks (§2.3). All models and experiments are implemented using PyTorch and are publicly available under https://github.com/gdewael/maldi-nn. The following sections outline each component of our study in detail.

### 2.1. Data

A first dataset is the recently published DRIAMS database (Weis et al., 2022). It is used to both pretrain Maldi Transformer and to fine-tune it on antimicrobial resistance (AMR) prediction (binary classification using a dual-branch recommender system). DRI-AMS contains a total of 250 070 spectra, originating from four hospitals in Switzerland. For AMR prediction, the same data splits are used as in earlier work (De Waele et al., 2023). Briefly, DRIAMS-A spectra from before 2018 are split to the training fraction, whereas DRIAMS-A spectra measured during 2018 are evenly split between validation and test set. For pre-training, the same splits are used, but all spectra from DRIAMS-B, -C, and -D are additionally added to the training set. Apart from AMR measurements, DRIAMS contains species labels for many spectra. These labels are derived from the species identification pipelines included with the MALDI-TOF MS machines. After processing (see Appendix A), the pre-training DRIAMS dataset spans 469 species.

The second dataset consists of the public Robert Koch-Institute (RKI) database (Lasch et al., 2023). The final used dataset is a private historical database containing more than 2400 taxonomic reference strains, cultured and analyzed by the Laboratory of Microbiology at Ghent University. In this manuscript, both datasets will be abbreviated as RKI and LM-UGent, respectively. The RKI dataset contains MALDI-TOF mass spectra from highly pathogenic bacteria, covering similar species as in DRIAMS. The LM-UGent dataset, on the other hand, includes a broader taxonomic range. Both datasets are used for fine-tuning on species identification (multi-class classification). As in Mortier et al. (2021), in order to create a challenging training-validation-test split, spectra are split in such a way that there is no overlap in terms of strains. As a consequence, the models are tested whether they can identify unseen strains of (seen) species. The following rules are used in data splitting: all spectra for a species are assigned to the training set if that species only has one strain in the dataset. If the species has more than one strain, strains are split such that 80% of strains of that species are assigned to the training set, and the other 20% evenly split between validation and test set (with a floor value of at least one strain being assigned to either validation or test set). The total number of species in the RKI training set spans 270, of which 106 and 108 are present in the validation and test set, respectively. For the LM-UGent dataset, these numbers are 1088, 200, and 202, for the training, validation and test set, respectively. For more details on the LM-UGent dataset, the reader is referred to Mortier et al. (2021). Table 1 lists a summary of the sizes of all used data.

**Table 1.**
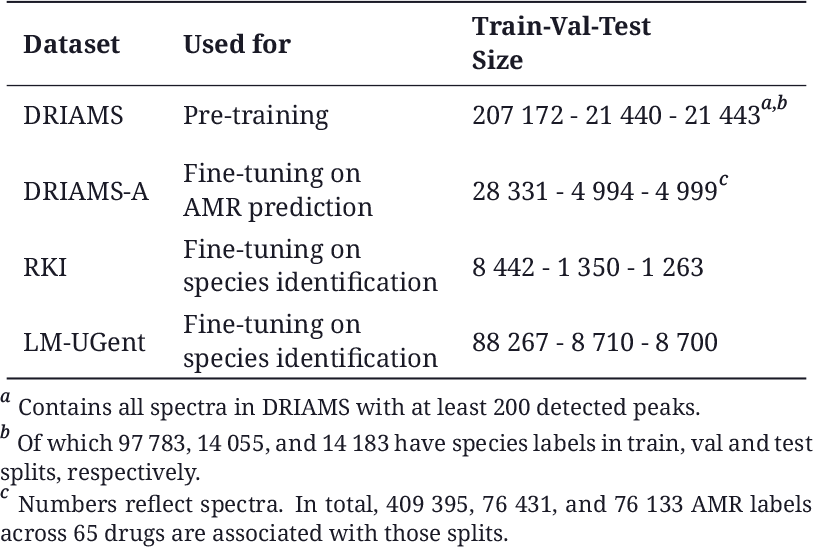
Data sources used, along with their use and sizes (in number of spectra).

All MALDI-TOF mass spectra are preprocessed using standard practices (Gibb and Strimmer, 2012). Following Weis et al. (2022), all spectra undergo the following steps: (1) square-root transformation of the intensities, (2) smoothing using a Savitzky-Golay filter with half-window size of 10, (3) baseline correction using 20 iterations of the SNIP algorithm, (4) trimming to the 2000-20000 Da range, (5) intensity calibration so that the total intensity sums to 1. For Maldi Transformer, peaks are then detected on the preprocessed spectrum using the persistence transformation, introduced by Weis et al. (2020b). While the original publication proposes this algorithm to nullify other preprocessing steps, we find that prior preprocessing steps help peak detection (see Appendix B). As in Weis et al. (2020b), we select the highest 200 peaks for every spectrum as inputs^1^. Maldi Transformer is compared against baseline methods, which require a fixed-length input. For these models, instead of running peak detection as a last preprocessing step, spectra are instead binned to a 6000-dimensional vector by summing together intensities in intervals of 3 Da.

### 2.2. Maldi Transformer

#### Model

To introduce Maldi Transformer, let us denote a spectrum as a set of *m* peaks 𝒮 = 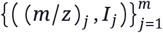 with each peak characterized by its (*m*/*z*) value and (preprocessed) intensity *I*. An annotated dataset 𝒟 then consists of a set of *n* samples 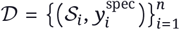, with *y*^spec^ the species label.

Maldi Transformer requires a single input representation ***x*** ∈ ℝ^*d*^ for each peak ((*m*/*z*) _*j*_, *I* _*j*_). To achieve this, intensities *I* are linearly transformed to a *d*-dimensional space. Similarly, (*m*/*z*) values are embedded to sinusoidal positional encodings (Vaswani et al., 2017). For any *m z* value, the positional encoding (*PE*) at feature index *f* is

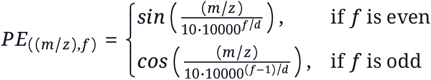

Note that this formulation is identical to the one described in Vaswani et al. (2017), apart from the factor 10 division of the (*m*/*z*) positions. This division is performed to bring the numerical range of (*m*/*z*) values (2000 Da - 20000 Da) closer to the numerical range of positional indices for which this equation was originally designed^2^.

The sinusoidal (*m*/*z*) embedding and linear intensity *I* embedding are summed to a single input representation per peak ***x*** _*j*∈ {1,…,*m*}_. A trainable [CLS] vector is prepended to the input for spectrum-level prediction, following common practice (Devlin et al., 2018; Dosovitskiy et al., 2020). The final input to the encoder-only transformer is, hence, ***X*** ∈ ℝ^201×*d*^, with 201 the number of peaks plus the [CLS] token, and *d* the hidden dimensionality of the model. The encoder-only transformer processes ***X*** to a spectrum-level output embedding ***p*** _[CLS]_ and output peak-level embeddings ***p*** _*j*∈ {1,…,*m*}_. The design of the transformer encoder blocks follows current state-of-the-art practices (Appendix E Figure 6) (Narang et al., 2021). An overview of the model is visualized in Figure 1.

**Figure 1.**
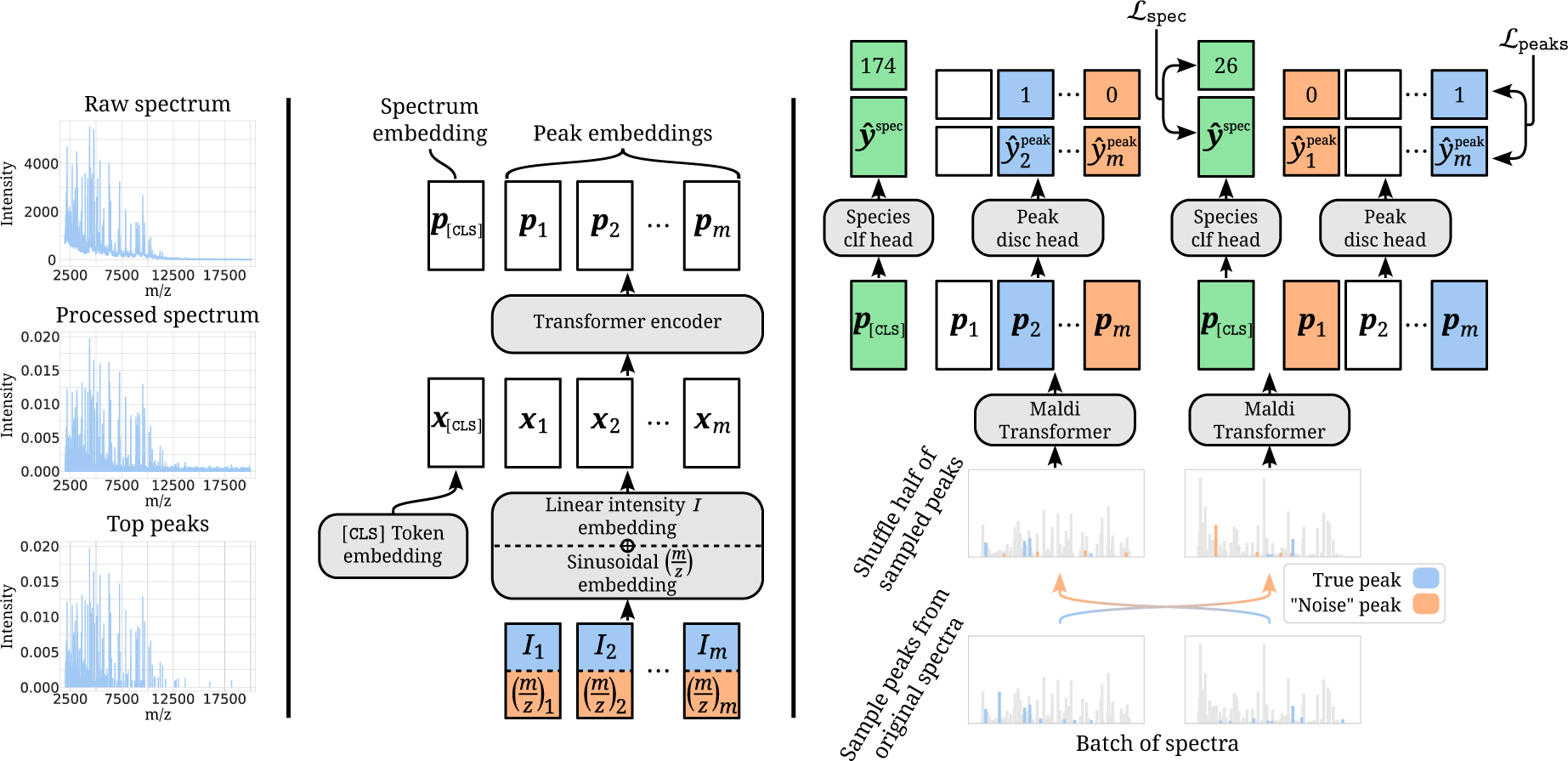
**Left:** Representation of MALDI-TOF mass spectra. A raw spectrum is preprocessed, after which a topological persistence transformation is performed to detect peaks, resulting in a sparse representation of the spectrum (Weis et al., 2020b). **Middle:** Maldi Transformer. A peak is characterized by an intensity *I* and (*m*/*z*) value. Both are embedded to higher dimensional space by a linear layer and sinusoidal embedding, respectively, and are then summed. A `[CLS]` token is prepended and the resulting vectors ***x*** are sent through multiple Transformer encoder layers. The resulting output vectors ***p*** _*j*∈ {1,…,*m*}_ can be used as peak embeddings. Additionally, ***p***_[CLS]_ can be used as a summary embedding of the whole spectrum. **Right:** Proposed peak discrimination pre-training strategy. In a mini-batch of spectra, 15% of peaks are randomly sampled, half of which are shuffled among all spectra in the batch. The resulting spectra are encoded with Maldi Transformer. The resulting peak embeddings ***p*** _*j*∈ {1,…,*m*}_ are sent through a linear layer (“Peak disc head”) trying to distinguish original peaks (blue) from shuffled “noise” peaks (orange). Spectrum embeddings ***p***_[CLS]_ are sent through a separate linear output layer (“Species clf head”) to predict species identity (green). The numbers 174 and 26 refer to example ground-truth class indices *c* = *y*^spec^ (indicating the microbial species of the spectrum).

#### Pre-training task design

To boost the performance of Maldi Transformer on supervised tasks with limited labeled data, a novel self-supervised pre-training strategy is designed. We propose to pre-train Maldi Transformers as peak discriminators. That is, in a batch of spectra, some peaks are randomly sampled to use for training. Following Devlin et al. (2018), we sample 15% of the peaks. Half of those sampled peaks are shuffled among all spectra, while the others are kept as part of their original spectrum. Using the shuffled peaks as negative “noise” peaks in a spectrum, a discriminative model is trained to distinguish the noise peaks from the sampled original ones using the cross-entropy loss.

More formally, for a spectrum 𝒮, let us denote its version with shuffled peaks as 𝒮^shuff^. Further, let ***j***_train_ = {*j*_1_, …, *j*_*s*_} denote the indices of its sampled peaks. Maldi Transformer processes 𝒮^shuff^ to output peak-level embeddings ***p*** _*j*∈{1,…,*m*}_, which are then processed to predictions 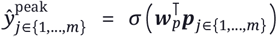, with ***w***_*p*_ a learnable weight vector. The peak discrimination loss is, then, given by

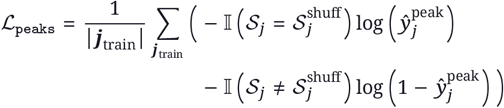

with 𝕀 the indicator function, and *j* indexing single peaks from the spectrum 𝒮

Complementary to the peak discrimination strategy, the spectrum embedding ***p*** _[CLS]_ is sent to a multi-class linear output head to predict the microbial species identity *y*^spec^ of the original spectrum^3^: 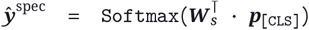, with ***W***_*s*_ a learnable weight matrix. Species predictions ***ŷ*** ^spec^ are optimized with multi-class cross-entropy: 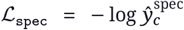, with *c* = *y*^spec^ the ground-truth class index. The final pre-training loss function is the sum of peak discrimination binary cross-entropy and species identification multi-class cross-entropy: ℒ = ℒ_peaks_ + *λ* ℒ_spec_, with *λ* ∼ Bern 0.01. The species identification loss ℒ_spec_ is only randomly applied in 1% of the training steps. This is performed because, empirically, overfitting of the species classification task is observed when applied at every training step (see Appendix E Figure 7). A visual representation of the entire pre-training approach is shown in Figure 1. A more-thorough description of the whole pre-training strategy can be found in Appendix C Algorithm 1.

Our novel peak discrimination pre-training strategy is proposed due to conceptual issues with porting established pre-training approaches to the MALDI-TOF MS domain, a point further elaborated on in Appendix C. We benchmark this novel pre-training strategy against alternatives in §3.2.

#### Model configurations

We train Maldi Transformer in four different sizes: S, M, L, and XL. Model sizes are chosen so that the total weight numbers roughly correspond to the ones in De Waele et al. (2023). Table 2 lists the size of all models, along with some hyperparameter settings. The Adam optimizer is used to pretrain all models in BFloat16 mixed precision (Kingma and Ba, 2014). Gradients are clipped to a norm of 1. A batch size of 1024 is applied for all models. A linear learning rate warm-up is applied over the first 2500 steps, after which the learning rate remains constant (Table 2). During training, a dropout of 0.2 is applied in the GLU feedforward and over the attention matrix.

**Table 2.** Pre-training configurations for different Maldi Transformer sizes. Species clf *λ* refers to the probability that the species identification loss is applied per training step.

| Hyperparameter | Model |  |  |  |
| --- | --- | --- | --- | --- |
|  | S | M | L | XL |
| # params | 1.65M | 3.27M | 6.92M | 14.84M |
| # layers | 4 | 6 | 8 | 10 |
| hidden dim $d$ | 160 | 184 | 232 | 304 |
| # heads | 8 | 8 | 8 | 8 |
| Learning rate | 5e-4 | 5e-4 | 5e-4 | 3e-4 |
| Pre-training steps | 500 000 | 500 000 | 500 000 | 400 000 |
| Species clf $\lambda$ | Bern(0.01) | Bern(0.01) | Bern(0.01) | Bern(0.005) |

### 2.3. Experimental set-up of downstream tasks

As mentioned in §2.1, Maldi Transformer’s performance is validated on three downstream supervised tasks. For all three tasks, the pre-trained model is plugged in at initialization and all weights are finetuned (i.e. no weight freezing). A task-specific linear output head ***W***_*out*_ projecting the spectrum embedding ***p*** _[CLS]_ to the desired output space is trained from scratch. For AMR prediction, Maldi Transformer is used as a spectrum embedder in a dual-branch neural network recommender system. To compare its performance against previous results, recommenders are trained with the four best-scoring drug embedders in De Waele et al. (2023). To benchmark Maldi Transformer in terms of species identification, the RKI and LM-UGent datasets are used. Species identification is compared to MLP baselines, Logistic Regression, Random Forest, and k-nearest neighbors (k-NN) models. Details on the exact fine-tuning setups for Maldi Transformer, as well as training details for all baselines are found in Appendix D.

The LM-UGent dataset covers a broader species diversity than the clinical species found in the pretraining set. As such, this dataset can be considered out-of-distribution for the pre-trained Maldi Transformer. For this reason, a domain adaptation step on the pre-trained Maldi Transformer is performed before supervised fine-tuning. The domain adaptation step consists of pre-training Maldi Transformer for 20 000 additional steps in the same fashion, but now using the LM-UGent dataset, instead of DRIAMS^4^.

## 3. Results

In the following sections, we first examine Maldi Transformer’s predictive performance compared to baselines (§3.1). We experimentally validate the soundness of our model design through ablations (§3.2). Finally, we exploit Maldi Transformer’s denoising properties in §3.3.

### 3.1. Maldi Transformer improves performance on downstream tasks

Pre-training curves for Maldi Transformer are shown in Appendix E Figure 8. Maldi Transformer spectrum embeddings after pre-training are visualized in Appendix E Figure 9. After pre-training, Maldi Transformer is fine-tuned w.r.t. a downstream task. Maldi Transformer’s downstream performance is compared to preprocessed and binned baselines. For AMR prediction, it is compared to MLPs of similar sizes. For species identification, it is additionally compared to non-neural network baselines.

Figure 2 shows the experimental results for all downstream tasks. For AMR prediction, models are evaluated in terms of their (per-spectrum-average) ROC-AUC and Precision@1 of the negative class (Prec@1(-))^5^. Appendix E Figure 10 additionally shows their performance in terms of (drug) macro ROC-AUC. For species identification, models are evaluated in terms of species- and genus-level accuracy. In general, Maldi Transformer obtains superior performance in comparison to other tested methods. For AMR prediction, Maldi Transformer consistently outperforms all MLP models in terms of ROC-AUC. In terms of the Prec@1(-), Maldi Transformer is sometimes outperformed by MLP models, but the best-scoring model overall is still one using Maldi Transformer (i.e. the XL model paired with a Morgan Fingerprint drug embedder). On drug Macro ROC-AUC (Appendix E Figure 10), Maldi Transformer consistently outperforms MLPs. As this metric computes how well the model separates spectra in terms of their resistance status per drug, this metric is especially apt for showing the advantage of pre-training to improve spectral representations.

**Figure 2.**
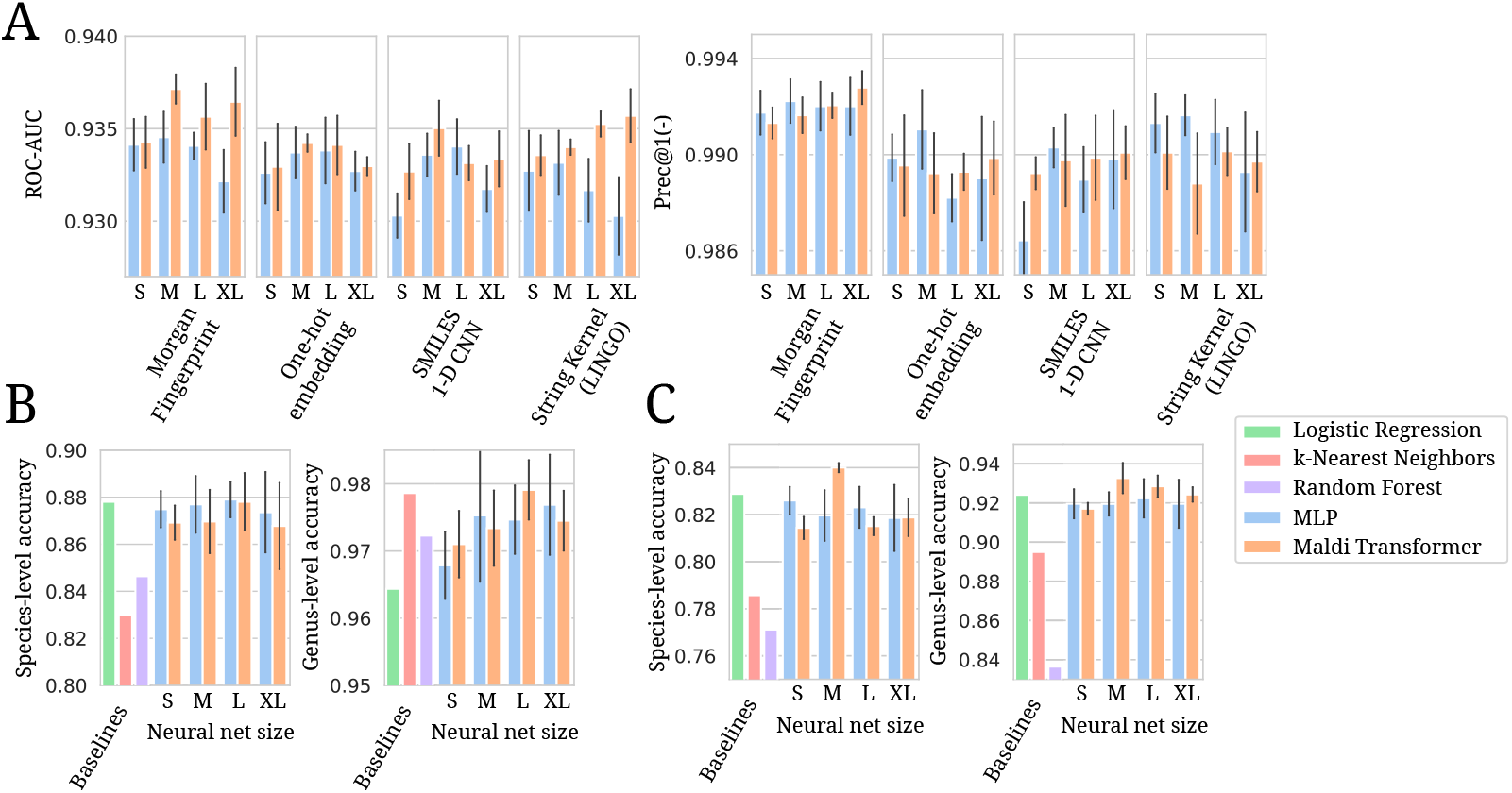
Barplots of all main experimental results. Error-bars indicate standard deviation over five independent runs. Neural network model sizes range from S to XL. Details per model size for Maldi Transformer and MLP models are listed in Table 2 and Appendix D Table 3, respectively. **A:** AMR prediction results on DRI-AMS. Performances are shown for models using four different drug embedders in a dual-branch recommender system. **B:** Species identification results on the RKI dataset. **C:** Species identification results on the LM-UGent dataset.

For species identification on the RKI dataset, Maldi Transformer does not always provide a clear advantage in terms of species-level accuracy. For genus-level accuracy, however, the large Maldi Transformer variant achieves the best performance, with a k-NN model providing competitive results.

On the larger and more-difficult LM-UGent dataset, results are again more convincing. The best-performing species-level model is the medium-sized Maldi Transformer, convincingly beating other modsels with 84% accuracy. The same results are obtained on genus-level accuracy, where Maldi Transformer consistently beats other models.

In order to gain a better understanding into when Maldi Transformer might provide advantages over other models, an experiment is run on the LM-UGent dataset. The experiment entails progressively leaving out species (classes), keeping only the top-*n* most occurring species, and training and evaluating models on that subset. The results, visualized in Appendix E Figure 12, show that by gradually decreasing the number of classes, and, hence, the difficulty of the prediction task, Maldi Transformer’s advantage on LM-UGent gradually disappears.

### 3.2. Maldi Transformer pre-training ablation

To further validate the efficacy of the proposed Maldi Transformer, its performance is compared against four alternative realizations of transformers on MALDI-TOF MS data (i.e. using four alternative pre-training strategies). The first takes inspiration from the masked language model (MLM) BERT (Devlin et al., 2018). Here, intensity values of peaks are randomly masked out, and a transformer is pre-trained to predict the original values back using the mean-squared error loss. In the second, the same procedure is followed, but instead of regressing masked intensities, on the output-level, intensities are binned into discrete height bins. A model is then pre-trained to classify the bin of the masked intensity using the cross-entropy loss. The third pretraining strategy consists of the same as the previous, but masking (*m*/*z*) values instead, much as like proposed in Bushuiev et al. (2024). In the last, a discriminative model much like our final proposed strategy is trained. The difference in this model is that negative peaks are sampled from some estimated distribution of peaks, instead of generating negative peaks by shuffling. All alternative pre-training strategies are described in greater detail in Appendix C.

In Figure 3.2A, it is observed that our proposed peak shuffling pre-training strategy consistently outperforms competing approaches. Our results show merit to the design of a pre-training task specifically adapted to the MS domain. In comparison, MLM-based designs ported from the language domain result in comparatively worse performance.

**Figure 3.**
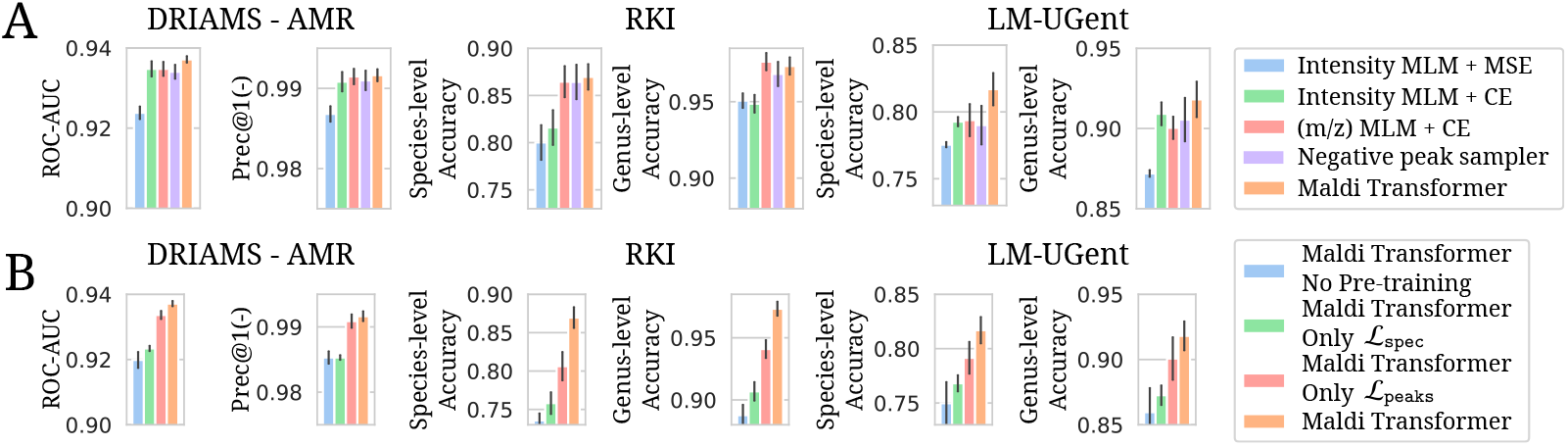
Barplots of all ablation results. Only results for Medium-sized transformers are shown. The final model as discussed in §2.2 is labeled as Maldi Transformer. Error-bars indicate standard deviation over five independent runs. For DRIAMS AMR prediction, only results using a Morgan Fingerprint drug embedder are shown. For the LM-UGent dataset, results are shown without the domain adaptation step, as this step was also not performed for ablation models. See Appendix E Figure 13 for the effects of this step. **A:** Testing different ways of pre-training a transformer on MALDI-TOF MS data. **B:** Leaving out different parts of the final pretraining strategy.

Figure 3.2B shows ablation results when leaving different parts out of the final pre-training strategy. Maldi Transformer is compared to models where one of the two loss components are left out: either the peak discrimination loss (ℒ_peaks_), or the species identification loss (ℒ_spec_). It is also compared to a transformer model without pre-training whatsoever. It can be seen that the latter strategy (i.e. training a separate transformer from scratch for every supervised task), delivers subpar performance. In addition, only pretraining on species identification does not help much. The biggest gain is made from the peak discrimination task. Yet, overall, the effects are additive, showing that each part contributes to the final performance.

A final ablation is to test the choice of peak detection. Here, the persistence transformation algorithm (Weis et al., 2020b) used in Maldi Transformer is tested against the established MALDIquant approach to detect peaks (Gibb and Strimmer, 2012). The results of this ablation are shown in Appendix E Figure 11.

### 3.3. Maldi Transformer model analysis

Due to its pre-training design, Maldi Transformer outputs the probability of a peak belonging to its spectrum for every peak. More formally, let us denote the output probability of a peak *j* belonging to its spectrum 𝒮 as 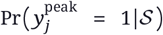. This quantity can be obtained through the pre-trained Maldi Transformer: 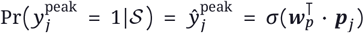. During pretraining, as peaks are shuffled between spectra, one expects to encounter both positives and negative true labels. In this section, we examine how peak output probabilities 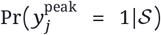 may be used during inference (i.e. without shuffling) using the medium-sized Maldi Transformer. The interpretation of probabilities requires a model to be calibrated. This property is further examined in Appendix E Figure 14. Results in the following paragraphs are derived from the DRIAMS pre-training test set.

#### Maldi Transformer denoises spectra

In Figure 4A, a randomly selected *Staphylococcus epidermidis*^6^ spectrum is visualized, each peak colored according to its output probability 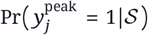. Most peaks are (correctly) assigned a high probability of belonging to their spectrum. The peaks with a low probability could be mistaken by the model, or could represent “true” noise in the spectrum. Such noise may still be present in the final spectrum due to, for example (1) noisy readouts from the spectrometer, or (2) shortcomings in preprocessing. In order to validate this hypothesis, it makes sense to look at patterns across multiple spectra. In Figure 4E, all *S. epidermidis* spectra are visualized together, each peak as a single dot. Black dots represent peaks that are predicted to be noise with high probability 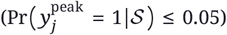. In the magnified parts of the plot, it can be seen that blue dots cluster together, meaning that *S. epidermidis* spectra often have peaks in the same places. Black dots mainly fall outside or on the edges of those clusters, signifying that those peaks are rightly picked up by the model as noise. Consequently, Maldi Transformers can serve a broader purpose as spectrum denoisers, in addition to their supervised learning capabilities.

**Figure 4.**
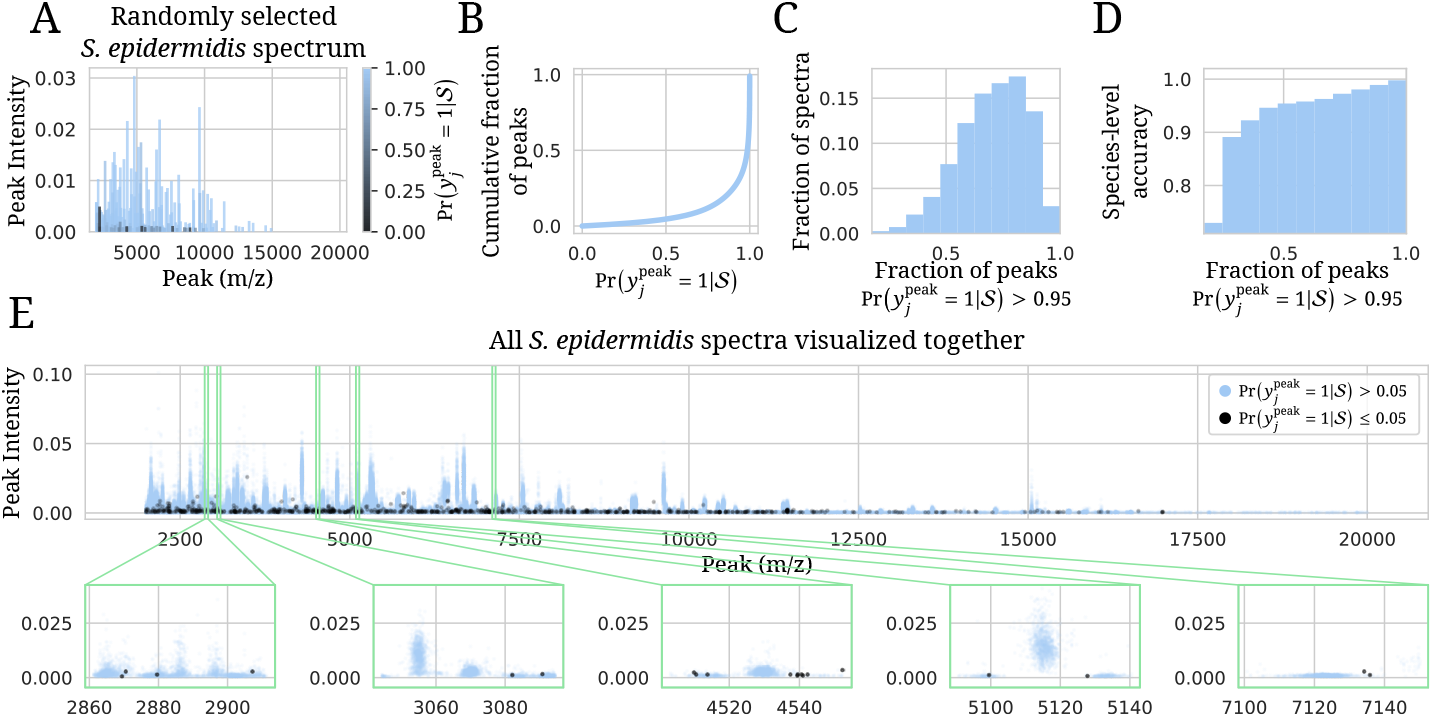
(Medium-sized) Maldi Transformer analysis on DRIAMS pre-training test data. Note that no shuffling of peaks across spectra is performed to generate these plots. Each spectrum contains their original detected peaks. **A:** A randomly selected spectrum, each peak colored by the model’s output probability of that peak belonging to that spectrum or not 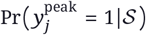. **B:** Empirical cumulative distribution of those output probabilities over all spectra in the DRIAMS test set. **C:** Histogram of fraction of peaks confidently (> 0.95) predicted as “true” peaks per spectrum. **D:** Species-level accuracy in function of fraction of peaks confidently (> 0.95) predicted as “true” peaks per spectrum. **E:** All *S. epidermidis* spectra visualized together, every peak as a separate dot. Peaks confidently predicted as “noise” (≤ 0.05) are shown in black. Zoomed in subplots show that “noise” peaks originate outside clusters of usual peak locations.

#### Noisy peaks are indicative of lower downstream performance

In order to better grasp the effect of noisy peaks on spectra, peak predictions are examined across all spectra in the DRIAMS test set. Figure 4B shows the empirical cumulative distribution of peak output probabilities. Only approximately 5% of all peaks are assigned a probability of belonging to their respective spectrum 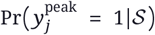 of lower than 50%. Conversely, half of the peaks are predicted with a probability of 98.5% or greater. This shows that while noisy peaks do exist, they typically make up a small percentage of the overall input spectrum.

Noisy peaks may not uniformly occur across the dataset. Some spectra may have more noisy peaks than others. A good spectrum quality statistic can, hence, be the fraction of peaks that are confidently predicted as “true”, e.g. with a probability greater than 95%. Figure 4C shows the distribution of spectra in function of this statistic. A slight left tail in the distribution signifies that some spectra have a large amount of noisy peaks. Aproximately 10% of the spectra have more than half of their peaks predicted with a probability smaller than 95%. Figure 4D shows that this statistic is also indicative of predictive performance. Spectra with more confidently “belonging” peaks have a higher species-level accuracy (on DRI-AMS test set labels). Species-level accuracy ramps up from 90% (or lower) for spectra with a lot of noisy peaks, up to nearly-perfect accuracy for spectra with (almost) all “predicted true” peaks^7^. The fraction of “predicted true” peaks also correlates with prediction certainty (see Appendix E Figure 15). These results showcase Maldi Transformer’s ability to not only improve performance, but also provide further insights into the data.

## 4. Discussion

Maldi Transformer adapts the transformer model for MALDI-TOF MS data, and proposes to pre-train said model using a novel peak shuffling approach. Using this novel self-supervised task design, we show state-of-the-art (or competitive) performance on the most important real-world use-cases of MALDI-TOF MS. Importantly, we show that our novel self-supervised pretraining task design delivers better performance compared to competing pre-training paradigms.

Experiments on increasingly simpler species identification tasks (Appendix E Figure 12) shows that the advantage of Maldi Transformer is most pronounced on more difficult tasks. This perhaps explains why experimental results on the smaller RKI dataset do not clearly show an advantage for Maldi Transformer. On the comparatively more complex tasks, such as 1088-way classification on LM-UGent and dual-branch recommendation on DRIAMS, Maldi Transformer consistently delivers state-of-the-art performance.

As with many subfields of machine learning, size, quality, and diversity of data constitute the biggest determinant of performance. In this study, an apparent ceiling in representational capacity has been obtained given the available public data. This is evidenced by the fact that embeddings of increasingly larger variants of Maldi Transformer do not uncover more finegrained clusters (Appendix E Figure 9). In addition, in fine-tuning, XL variants of Maldi Transformer typically do not give advantages over the M or L models. The size of the XL Maldi Transformer (∼15M weights), however, is still small by self-supervised transformer standards. We hypothesize that the representational capacity of larger transformers is simply not necessary for the relatively-simple datasets that are available in the MALDI-TOF MS domain. Because of this, we argue that more efforts to collect and publish data may be the most important factor for MALDI-TOF-based machine learning research to continue to flourish. Using more complex data, it is possible to formulate more complicated learning tasks, such as the recommender system in De Waele et al. (2023), that benefit more from the increased representational capacity of Maldi Transformer. As spectral data is routinely generated within hundreds, if not thousands, of hospitals, we envision collaboration efforts with healthcare (as in Weis et al. (2022)) to play a crucial role in continued data collection and publishing.

Deep learning models have been hailed as feature extractors. For this class of models, it is typically argued that it matters less how features are presented to the model, as they can compose the relevant features themselves in their hidden representations. We would caution against this perspective, and state that how inputs are presented to a deep learning model could have far-reaching impacts on what the model can learn. Taking language as an example, character-level language models traditionally underperform the same models trained on (sub)word-level. In this work, MALDI-TOF mass spectra are presented to the model by their peaks. Because of this, any preprocessing steps and algorithms to determine peaks play crucial roles in the final performance of the model. Here, we present a small ablation study (Appendix E Figure 11), but we identify further experimentation with peak detection and preprocessing algorithms as a promising future research direction.

Our novel peak shuffling pre-training paradigm outperforms MLM techniques that have been ported from language and have been proposed in the metabolomics MS data domain (Bushuiev et al., 2024). MLM pre-training introduces noise in the spectrum by obfuscation. We hypothesize that MLM pre-training does not match the performance of peak shuffling due to noisy optimization. Consider, for example, a masked (*m*/*z*) language model. Given an intensity and a masked (*m*/*z*) of a peak, multiple possible (*m*/*z*) locations might exhibit a peak with said intensity. As the ground truth pre-masking *m z* value only represents one of multiple possible ground truths, the model is penalized for all but one of many correct answers. For this reason, we argue that MLM pre-training is ill-suited to the MS domain. It is expected that our proposed pre-training task design could be useful in other mass spectral domains. For this reason, we hope that our ablations (§3.2) and discussion in Appendix C serve as useful guidelines for adapting this work to other mass spectral data modalities, generated with MS instruments using alternative ionisation and separation methods. One potential limitation of our method is that it only computes a loss for a certain percentage of peaks per spectrum, similarly to MLM. A line of further research could, therefore, be to adapt autoregressive pre-training to the spectral domain using domain knowledge.

As the pre-trained model can be interpreted as learning peak co-occurences, this property is examined in §3.3. There, it is shown that Maldi Transformer can be used to detect noisy peaks. The predicted absence of noisy peaks is found to correlate with higher predictive performance. These insights go beyond those offered by off-the-shelf machine learning techniques. Paired with its state-of-the-art (or competitive) performance results, it can be concluded that Maldi Transformer enhances what biological practitioners get out of their MALDI-TOF MS data.

## Acknowledgements

This work was supported by Research Foundation - Flanders (FWO) [PhD Fellowship fundamental research grant 1153024N to G.D.W.]. W.W. also received funding from the Flemish Government under the “Onderzoeksprogramma Artificiële Intelligentie (AI) Vlaanderen” Programme

## A. DRIAMS species processing

DRIAMS contains species labels for many spectra. These species labels are derived from the species identification pipelines included with the MALDI-TOF MS machines. As such, some preprocessing steps are taken to make the labels as presentable as possible to an ML model.

Firstly, labels in DRIAMS containing the string `“not reliable identification”` are integrally deleted. Second, many spectra are labeled as `“MIX!species”`, indicating that the spectrum potentially contains an impure mixture of species. As such, species labels for these spectra are also not used. Additionally, species labels that occur fewer than five times in the training set are removed. Finally, species labels occurring only in the validation or test set, but not in the training set, are similarly removed. After processing, DRIAMS contains 469 different species labels. Note that the previous steps involve removing of labels, not spectra themselves. Corresponding spectra are still kept in the dataset, albeit without a species label.

## B. On preprocessing and persistence transformation

While Weis et al. (2020b) argue that persistence transformation nullifies the need for the parameter-heavy chain of preprocessing steps, we advocate for the opposite. To support this stance, a simple exploratory visualization is made.

In Figure 5, all *Bacillus anthracis*^8^ spectra in the RKI training set are put through two preprocessing chains. The first only does (1) trimming to the 2000-20000 Da range, (2) intensity calibration so that the total intensity sums to 1, and (3) persistence transformation, and keeps the 200 highest peaks. The second preprocessing chain performs all those steps preceded by variance stabilization, smoothing, and background removal, as in §2.1. Then, across all preprocessed spectra, the occurrence of detected peaks in bins of 3 Da is counted and plotted in Figure 5. The resulting profile can be considered a summary profile of all detected peaks in *Bacillus anthracis* spectra. The detected peaks with extra preprocessing result in a cleaner profile, with peaks more concentrated in regions that clearly correspond to biologically-informative signal (especially notable when inspecting the left tail of the spectra). In contrast, without prior preprocessing, detected peaks are seemingly more-randomly distributed, hinting that noise in the spectrum negatively affects peak detection.

**Figure 5.**
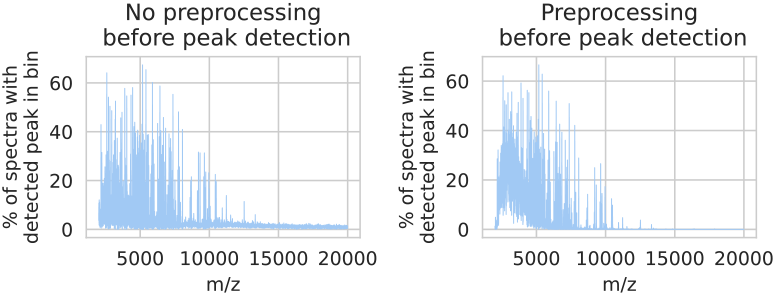
Visualization of summary profiles of *Bacillus anthracis* RKI training spectra obtained when doing peak detection with (R) or without (L) preprocessing first.

**Figure 6.**
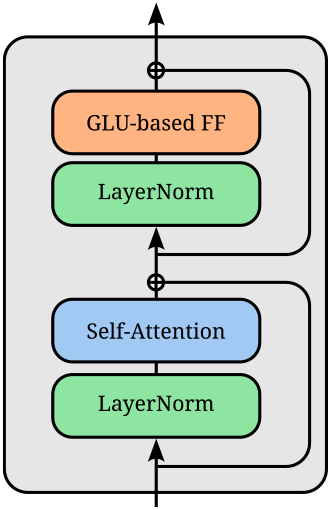
Utilized transformer encoder block. The network uses pre-LayerNorms and GeLU gated linear units (GLU) in the feedforward (FF) networks (Ba et al., 2016; Shazeer, 2020).

**Figure 7.**
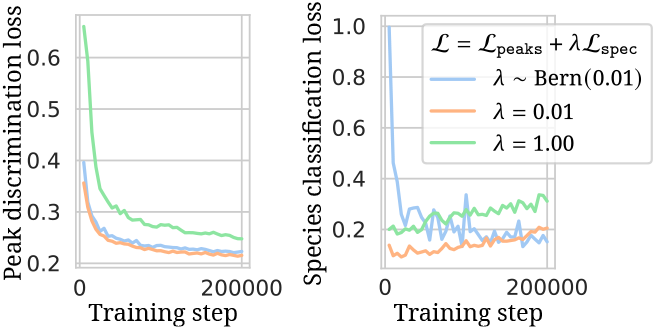
Pre-training dynamics using different strategies for combining the peak discrimination loss ℒ_peaks_ and the species identification loss ℒ_spec_. Loss on the validation set is shown. The figure is shown only for the first 200 000 pre-training steps using a medium-sized Maldi Transformer to illustrate. It is observed that if ℒ_spec_ is applied at every step (using a weight of 0.01 or 1.00), this component starts over-fitting well before the ℒ_peaks_ component is converged (training is performed for 500 000 steps total). By not applying ℒ_spec_ at every training step, the Adam optimizer momentum is dominated by ℒ_peaks_. As ℒ_spec_ is the “easier” task and more prone to overfitting, this regime benefits the final model.

**Figure 8.**
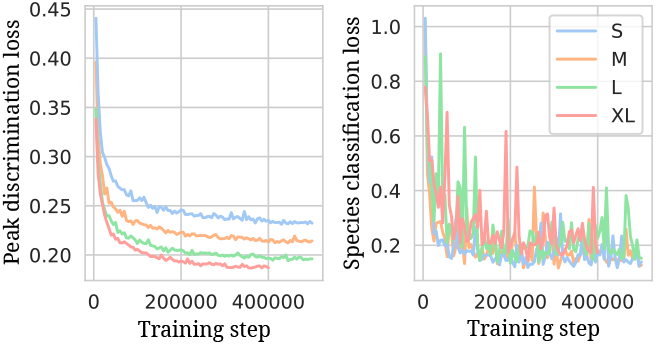
Pre-training validation loss curves for all Maldi Transformer model sizes.

**Figure 9.**
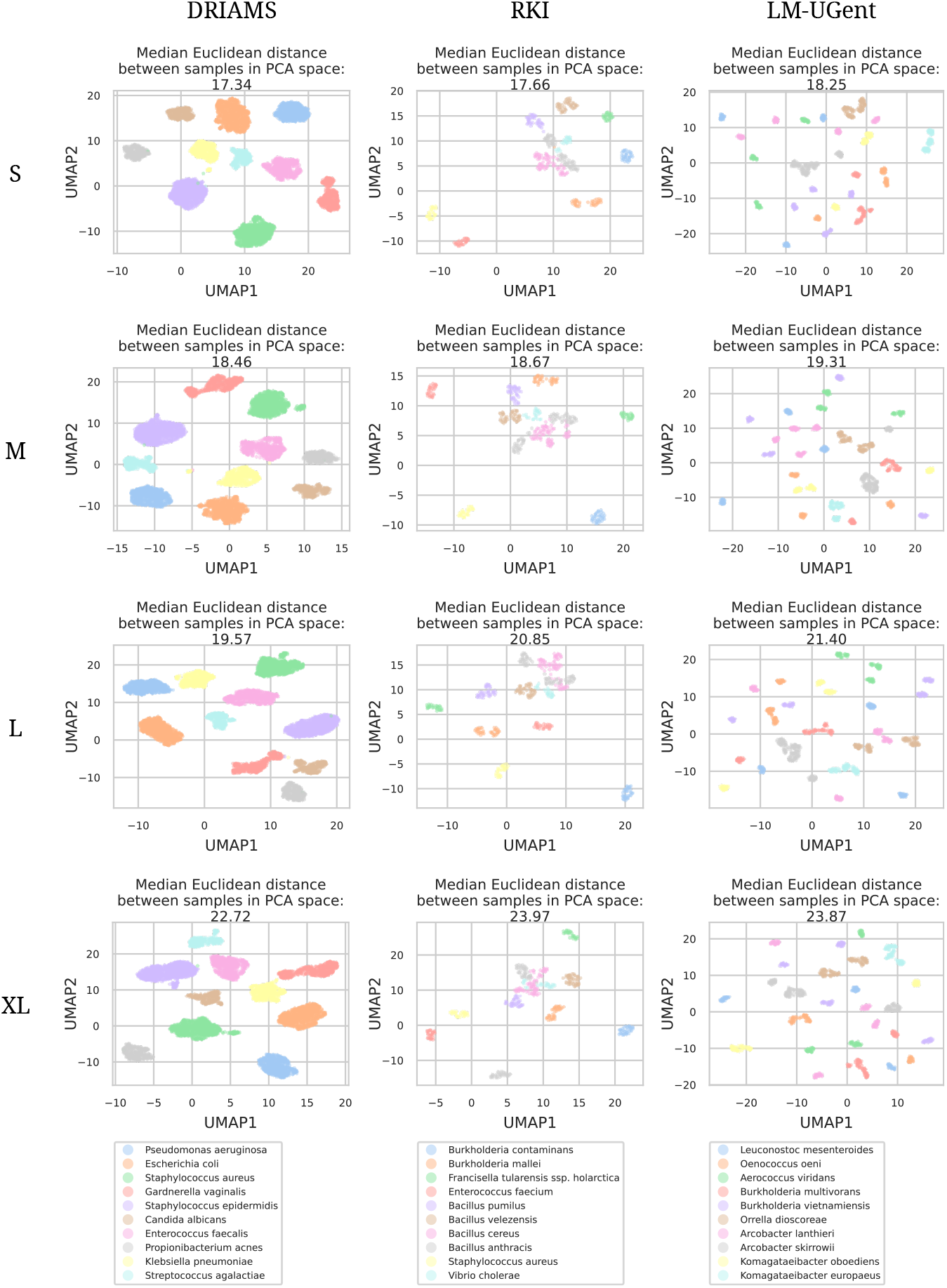
UMAP Embeddings of all validation spectra of all used datasets (columns), for every pre-trained (before fine-tuning) Maldi Transformer model size (rows). Only the 10 most occurring species in each dataset are used (legend). UMAP embeddings are obtained using ***p***_[CLS]_, standard normalized, then PCA transformed to 50 dimensions, then embedded in 2-dimensional space using UMAP. On top of every feature is the median euclidean distance between samples in the PCA space, before UMAP. It can be observed that, despite normalization and transforming to equal-dimensional spaces with PCA, the larger the pre-trained model, the more separated embeddings become. Notwithstanding, all models qualitatively find the same clustering.

**Figure 10.**
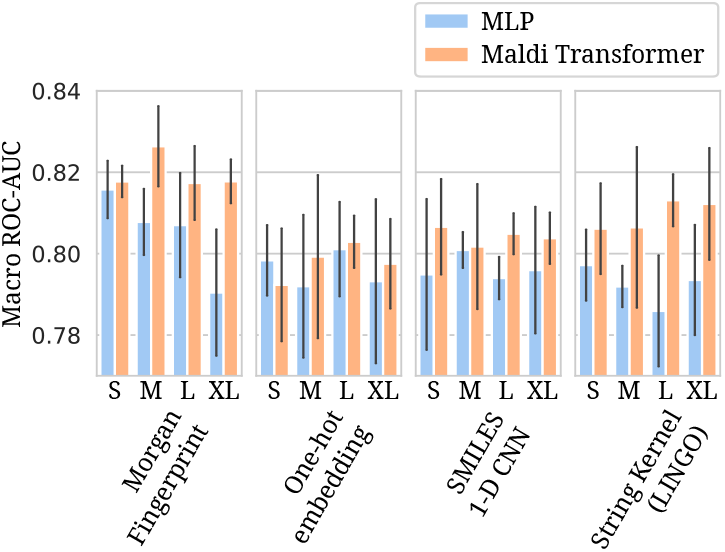
Barplots of AMR prediction on DRIAMS, evaluated by Macro (drug) ROC-AUC Error-bars indicate standard deviation over five independent runs. Neural network model sizes range from S to XL. Details per model size for Maldi Transformer and MLP models are listed in Table 2 and Appendix D Table 3, respectively.

**Figure 11.**
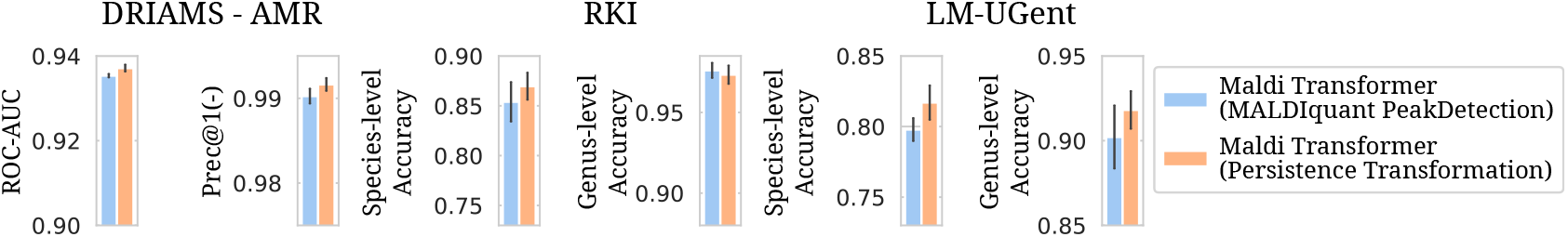
Barplots of all peak detection ablation results Only results for Medium-sized transformers are shown. The final model as discussed in §2.2 is labeled as Maldi Transformer (Presistence Transformation). The results for MALDIquant Peakdetection are obtained by pre-training and fine-tuning Maldi Transformer in exactly the same way, but with peaks detected using MALDIquant’s methodology (reproduced in Python). Error-bars indicate standard deviation over five independent runs. For DRIAMS AMR prediction, only results using a Morgan Fingerprint drug embedder are shown. For the LM-UGent dataset, results are shown without the domain adaptation step, as this step was also not performed for ablation models.

**Figure 12.**
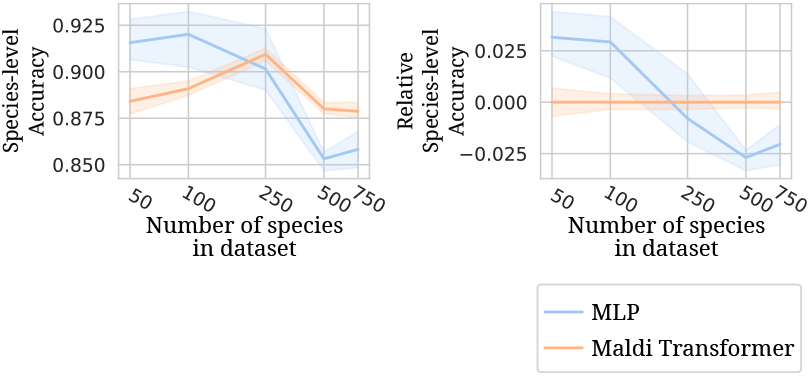
Lineplots of species-level accuracy on LM-UGent when trained and evaluated on a decreasing number of species. Species are filtered out on training data occurrency, always keeping the *n*-most-occurring species in the data. Errorband indicate standard deviation over three independent runs. The right hand plot shows the species-accuracy relative to the one obtained by Maldi Transformer.

**Figure 13.**
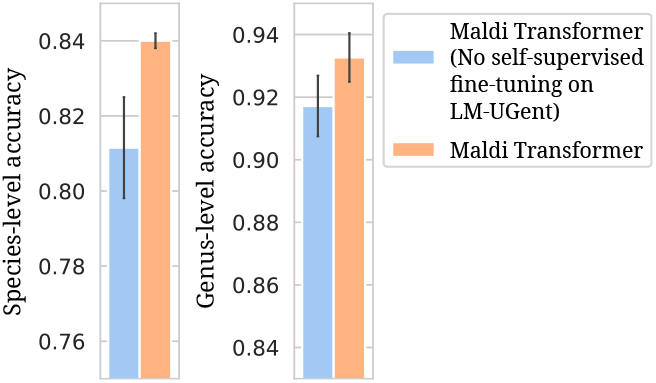
LM-UGent performance improves when, prior to supervised fine-tuning, the pre-trained model is first fine-tuned using the self-supervised training task. In the main text, we refer to this step as the domain adaptation step.

**Figure 14.**
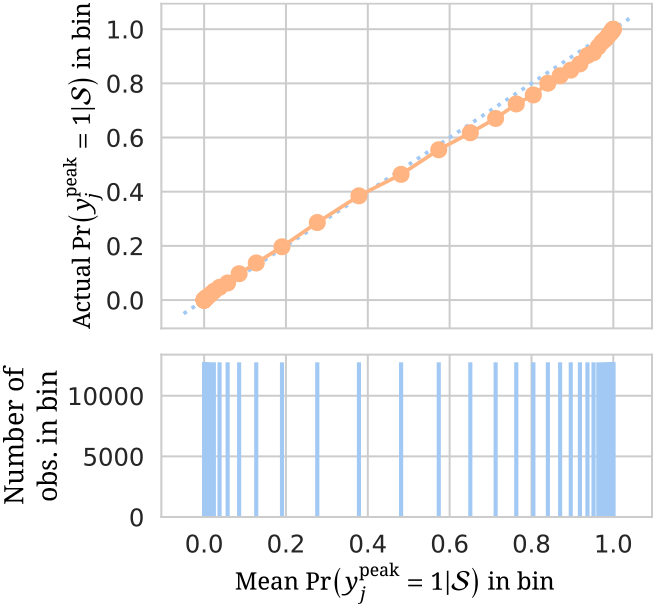
Calibration curve for pre-training peak discrimination using the medium-sized Maldi Transformer. On the DRIAMS test-set, the usual peak shuffling and peak discrimination is performed (see §2.2). Predictions are then sorted and split into equal frequency bins (on the x-axis). Within those bins, the average predicted value is plotted against the actual fraction of positive true labels in that bin. A calibrated model requires predictions to be interpretable in a frequentist manner, i.e. a sample with an output probability of 80% is expected to be positive 80% of the times. Hence, the aforementioned plot is expected to follow the diagonal. The plot shows that the model is reasonably-well calibrated, with slight overconfidence.

**Figure 15.**
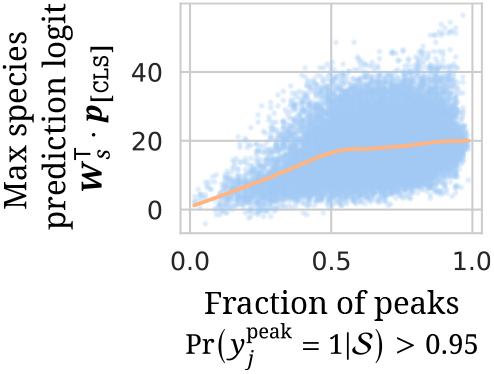
Max species prediction logit 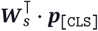 in function of number of confidently “belonging” peaks for a spectrum for DRIAMS test set spectra using the medium-sized Maldi Transformer. A higher max logit for the species prediction task corresponds to a higher max output probability post-softmax, and, hence, higher confidence. It is displayed that the model is more confident in its prediction for spectra with a higher number of confidently belonging peaks.

## C. Design of a self-supervised pretraining task

A large part of this study concerns the design of a self-supervised pre-training task for MALDI-TOF MS data. This is motivated by the scarcity of data in this domain. Through self-supervised learning, a greater representational capacity can be obtained, maximally utilizing patterns in the data. The following paragraphs chronicle thoughts and experiments w.r.t. the design of the SSL task.

MALDI-TOF MS spectra can be represented as sets of peaks. As each peak is characterized by an intensity and an (*m*/*z*) value, a natural parallel is drawn with language data. Instead of a sequence of words, a sequence of peaks is processed. A crucial difference is that, for language transformers, positional indices for each text “token” are set to the integer range numbering 0 to *n* − 1, with *n* the number of tokens in the input sequence. With MALDI-TOF MS spectra, positional indices (i.e. (*m*/*z*) values) are irregularly spaced and real-valued. Additionally, whereas words have a categorical identity in text, MALDI-TOF MS intensity is also real-valued. These two factors have to be taken into consideration when porting self-supervised learning techniques from one modality to the mass spectral domain.

In language transformer pre-training, perhaps the two most-classically quoted techniques are masked language modeling (MLM) (Devlin et al., 2018) and autoregressive modeling (AR) (Radford et al., 2018). The GPT series of models best exemplifies the success of the latter category (Brown et al., 2020). For MALDI-TOF MS, however, it is unclear how to autoregressively model a set of peaks. Autoregressive models imply a certain ordering of data, whereas this perspective is ill-fitting for the set-valued peaks input. For example, how does one train to generate the “next” peak given the previous peaks, if it is unclear how to define what the “next” peak is? For example, does one choose to order peaks by their height, or by their (*m*/*z*) value? If the model predicts the second-next peak instead of the next, is that necessarily wrong? If not, how to efficienctly construct the loss to take this into account?

Instead of answering the issues with autoregressively modeling a set-valued input, we may consider the alternative strategy: MLM, introduced by BERT (Devlin et al., 2018). For example, instead of masking out words, intensities can be masked, and a model can be trained to recover the intensities with the use of the mean-square error loss. This brings us to first ablated pre-training technique in §3.2. The training for this strategy is performed identical to the final strategy outlined in §2.2. The only difference being that the peak discrimination loss ℒ_peaks_ is exchanged for the mean-square error loss on masked peaks. Peaks to train on are similarly selected with 15% probability. Of those 15%, 80% are assigned masked intensities and 20% are left unchanged.

After fitting a regression MLM on intensities with limited success, a logical next step is to design a self-supervised classification task. A first choice may be to bin intensity values, and instead of regressing the masked intensities, classifying the intensity bin of the masked-out peaks. Here, we bin the intensity into 10000 possibe bins, and pre-train using a 10000-way classification task. A different route is to mask out the a MLM to classify the (*m*/*z*) bin of the masked-out (*m*/*z*) values of the peaks instead, and similarly train peaks. This approach has been previously described in (Bushuiev et al., 2024). Here, we bin the 2000-20000Da (*m*/*z*) -range into 1Da bins, and pre-train using a 18000-way classification task. Both methods are identically trained as the Intensity Regression MLM, described in the previous paragraph, but then using multi-class cross-entropy loss functions.

While MLMs on binned values already improve downstream performance results, some conceptual issues arise w.r.t. their application in the mass spectral data domain. Firstly, the masking procedure introduces a [MASK] token, which is only encountered during pre-training and fine-tuning. Hence, supervised fine-tuning is hampered due to a data distribution shift. Secondly, given a masked value of either intensity or (*m*/*z*), multiple possible answers may be feasible. For example, consider a masked (*m*/*z*) LM trained to predict the (*m*/*z*) of a peak given its intensity. Multiple locations for this peak may be feasible, but the model will penalize all but one. Hence, the plurality of ground-truth is not well-represented in the loss, and model pre-training is hampered. In order to overcome these issues, alternative pre-training techniques, taking inspiration from contrastive learning (Liu et al., 2021), are proposed. The following paragraphs describe the exact procedure resulting in the negative peak sampler model in §3.2.

Akin to contrastive learning, a per-peak classification task can ask a model whether a peak belongs to the rest of the spectrum or not. Just as in contrastive learning, this strategy requires to sample negative peaks to deliver negative samples. One way to generate negative peaks is to randomly generate them from some estimated distribution of peaks. Here, we estimate the distribution of peaks in two steps. First, all peak locations in the training dataset are collected: 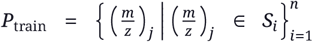, and a probability mass function over discrete bins is calculated:

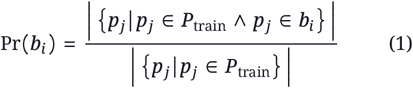

with bins chosen by 1 Da intervals: *b*_*i*_ ∈]*i, i* + 1], for *i* ∈ {2000, 2001, …, 19 999}. In other words, a probability mass function over 1 Da bins is created by counting how many times peaks fall into each bin in the training data. Next, within each bin, quantiles of the found intensities therein are calculated: 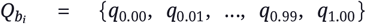. To sample a negative peak, an (*m*/*z*) value is first sampled by uniformly sampling a location within a sampled bin: (*m*/*z*) ∼ Pr *b*_*i*_ + Unif 0, 1. After, an intensity is drawn by uniformly sampling within a uniformly sampled interquantile range: 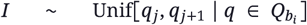, with *j* ∼ Unif{0.00, 0.01, 0.99}. The training for this strategy is performed identical to the final strategy outlined in §2.2. The only difference being that, instead of shuffling peaks around to generate negatives, negative peaks are now sampled. In every training step, 15% of the peaks are selected for training, half of which are exchanged for sampled negative ones.

As discussed in §3.2, generating negative peaks is not ideal for forcing the model to reason over peak cooccurrences. The model is allowed to learn any mis-representation in negatives as a shortcut. A way to circumvent the issues with generating negative peaks, is to use real ones. One way to present real peaks as negatives, is to take them from other spectra. This is where we land on the final self-supervised strategy that we test, and ultimately propose in our main text (see §2.2, Figure 1, and Algorithm 1).

### Algorithm 1

Maldi Transformer peak discrimination pre-training.

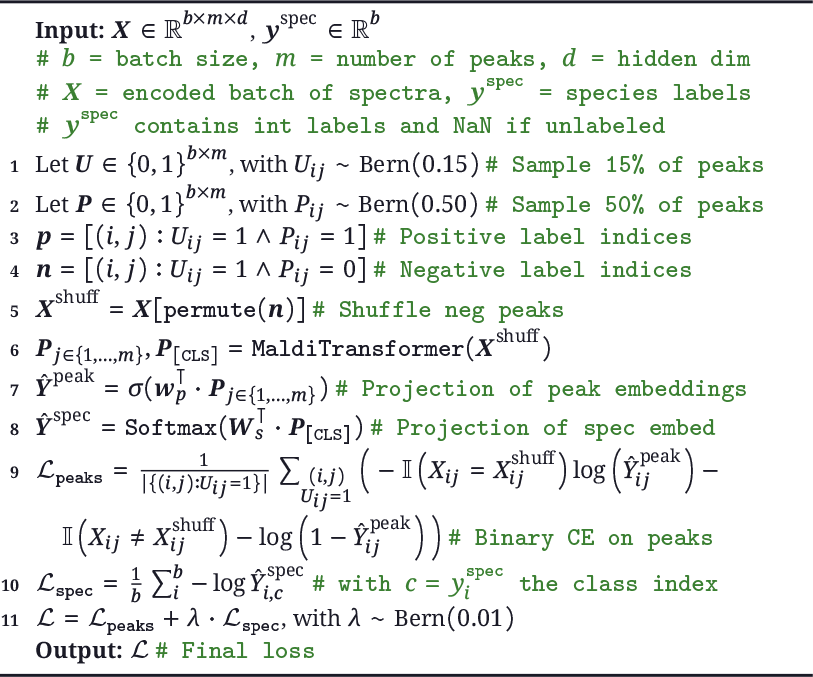

## D. Downstream tasks and baselines

For all three downstream tasks, the pre-trained model is plugged in at initialization and all weights are fine-tuned (i.e. no weight freezing). A task-specific linear output head ***W*** _*out*_ projecting the spectrum embedding ***p***_[CLS]_ to the desired output space is trained from scratch. All downstream models are similarly trained with the Adam optimizer with a batch size of 128 and dropout of 0.2. A linear learning rate warm-up over the first 250 steps is applied, after which the rate is kept constant.

For AMR prediction, training of Maldi Transformer-based recommenders is performed identical to the MLP-based baselines in De Waele et al. (2023). Briefly explained, for every combination of spectrum embedder (four sizes: S, M, L, and XL) and drug embedder (four types), six different learning rates ({1e-5, 5e-5, 1e-4, 5e-4, 1e-3, 5e-3}) are tested. For all these different combinations, five models are trained (using different random seeds for model initialization and batching of data). For every spectrum and drug embedder combination, only results from the best learning rate are presented; that is, the learning rate resulting in the best average validation ROC-AUC for that combination. The validation set is checked every tenth of an epoch. Models are trained for a maximum of 50 epochs, and their training is halted early when validation ROC-AUC hasn’t improved for 10 validation set checks. The checkpoint of the best performing model (in terms of validation ROC-AUC) is used as the final model. The baselines for the AMR prediction task are the models described in De Waele et al. (2023), which describes the recommender model structure in greater detail.

The fine-tuning of Maldi Transformer for species identification is performed in a similar way. The differences consist of: (1) ***W*** _*out*_ returning 270, or 1088, for the RKI and LM-UGent dataset, respectively, (2) five different learning rates are tested: ({1e-5, 5e-5, 1e-4, 5e-4, 1e-3}), and (3) validation species-level accuracy is tracked to halt training early (for a maximum of 250 epochs, and it is only checked once per epoch). As species identification is a multi-class classification task, models are then optimized using a softmax operation, combined with the cross-entropy loss. Species identification is compared to MLP baselines, Logistic Regression, Random Forest, and k-nearest neighbors (k-NN) models. All of these baselines are trained on preprocessed and binned spectra (see §2.1).

The MLP baselines are identical in construction to the spectrum embedders in De Waele et al. (2023), but with *n*-dimensional outputs instead of 64, with *n* the number of classes (see Table 3). MLP baselines are trained using the same strategy as with Maldi Transformer downstream fine-tuning. That is, for all model sizes, five different learning rates ({1e-5, 5e-5, 1e-4, 5e-4, 1e-3}) are trained five-fold. Model results from the best learning rate (in terms of validation species-level accuracy) are presented. Model training halts before its maximum of 250 epochs if validation species-level accuracy hasn’t increased in 10 epochs, and the model with the best validation accuracy is saved.

**Table 3.** All tested model sizes for the MLP baseline. Hidden sizes represent the evolution of the hidden state dimensionality as it goes through the model, with every hyphen defining one fully connected layer. *n* represents the number of output nodes. *n* equals 64, 270, and 1088 for DRIAMS AMR prediction, species identification on RKI, and LM-UGent, respectively.

| Size | # Weights | Hidden sizes |
| --- | --- | --- |
| S | 1.58M | 6000-256-128-$n$ |
| M | 3.25M | 6000-512-256-128-$n$ |
| L | 6.85M | 6000-1024-512-256-128-$n$ |
| XL | 15.09M | 6000-2048-1024-512-256-128-$n$ |

For non-neural baseline classifiers (Random Forest, Logistic Regression, and k-NN), a grid-search is performed to find optimal hyperparameters. Given the non-stochastic nature of their implementations, only one model is trained after hyperparameter tuning and, hence, only one test performance is reported. The parameter grid for Random Forest consists of `{max_depth = [25,50,75,100], min_samples_split = [2,5,10], max_features = [10,25,50,100]}`. All random forests are trained with 200 trees. For Logistic Regression: `{standardscaling = [True,False], L2_norm = [1e3,1e2,10,1,0.1,1e-2,1e-3]}. And for k-NN: {standardscaling = [True,False], n_neighbors = [1,2,3,4,5,6,7,9,10,25]}`.

## E. Figures and tables supporting the methods and results sections

## Footnotes

1 Note that Maldi Transformer can, in principle, deal with variable number of peaks per spectrum.

2 Language transformers are often trained with maximum sequence lengths between 512 and 2048. (Beltagy et al., 2020)

3 While this is not a purely self-supervised training objective, we justify its use with two reasons. First, in language, pre-training with a sentence-level [CLS] task boosts downstream performance (Devlin et al., 2018). Second, due to the integrated nature of MS manufacturers’ species identification pipelines, bacterial species labels are relatively easy to come by. As such, no manual labeling is necessary to obtain these labels. Additionally, it has to be noted that, as not all spectra in DRIAMS carry a species label, the species classification loss is only calculated for labeled spectra.

4 By performing domain adaptation, in contrast to the other supervised tasks, the task-specific output head ***W*** _*out*_ does not need to be initialized from scratch anymore, but can be copied from the pre-trained model, as it already has the right dimensions.

5 The spectrum-macro ROC-AUC evaluates the average ranking quality of recommended drugs. The Precision at 1 of the negative class evaluates the proportion of top-recommended drugs that are correct. These metrics are described in more detail in De Waele et al. (2023).

6 *Staphylococcus epidermidis* is the most occurring species in the DRIAMS pre-training test set.

7 Note that DRIAMS species labels are produced from the MS manufacturers’ software and models. It can be expected that it is relatively easy for a model to reproduce the labels (predictions) from another model. Hence, the nearly-perfect species-level accuracy on DRIAMS is not unexpected.

8 *Bacillus anthracis* is chosen because it is the most occurring species in the RKI training dataset.

## Notes

### Competing Interest Statement

The authors have declared no competing interest.

### Summary of Updates

This version makes minor changes to the previous version. Some grammatical errors have been fixed.

